# Climate warming reduces the energetic advantage of long-distance migration in a migratory bird

**DOI:** 10.64898/2026.05.07.723401

**Authors:** Martins Briedis, Joanna Wong, Detlef Becker, Martin Schulze, Dirk Tolkmitt, Paul Dufour, Steffen Hahn

**Author notes:** Corresponding author’s.

## Abstract

Climate warming is driving shifts toward shorter migration distances in migratory birds, yet the energetic mechanisms underlying this transition remain unclear. To understand how migratory birds balance energetic costs of movement with wintering benefits requires quantifying their energy expenditure across the annual cycle. Here, we present a novel approach to reconstruct annual energy budgets of free-flying birds from multi-sensor geolocator data on atmospheric pressure, ambient light, and activity. We apply this method to tracking data from Eurasian Wrynecks *Jynx torquilla* breeding in Central Europe. Tracking revealed striking individual differences in migration distance ranging from short-distance migrations of approx. 2000-2500 km to southern Europe and northern Africa to long-distance migrations of >5000 km to sub-Saharan Africa, with the former assumed to be a recently evolved strategy. Estimated mean daily energy expenditure across all individuals was 73.1 kJ/day, however, it was dependent on migration distance: long-distance migrants displayed 21-26% lower annual energy requirements than shorter-distance migrants despite 2-3 times higher costs of migration. The lower energy budgets of long-distance migrants resulted primarily from low thermoregulation costs during the ca. 5-month stationary non-breeding period. However, long-distance migrants faced extreme physiological demands during migration, with daily energy expenditure exceeding 9.5 times BMR approaching the upper limits of animal metabolic performance. We show that since 1950, climate warming has progressively reduced winter thermoregulation demands disproportionately benefiting and potentially promoting shorter-distance strategies. These results reveal shifting energetic trade-offs between migration and overwintering under climate change, potentially driving the evolution of migration.

## Introduction

Bird migration is a remarkable adaptive strategy that enables birds to exploit seasonal peaks in resource availability and avoid unfavourable environmental conditions [1,2], yet it comes with significant risks and energetic costs. Beyond the physically-taxing flight [3], long-distance migrants also face risks of adverse weather conditions and low food availability, particularly during long-distance transits over ecological barriers [4]. Despite these challenges, long-distance migration enables birds to access wintering habitats that offer improved conditions with warmer climates and more abundant food resources, which can reduce daily energy expenditure during the non-breeding season and thus, improve within-winter survival [5–7]. Conversely, individuals that undertake shorter migrations and winter in higher latitudes may avoid risks and conserve energy during the migratory phase but face the challenge of wintering in harsher environments with lower food availability and greater thermoregulatory demands.

When individuals from the same breeding population vary widely in migration distance, it provides a naturally occurring experimental setup to study how different migration strategies shape energetic trade-offs across the annual cycle [7–10]. This trade-off between longer migration distances or harsher wintering conditions is crucial for understanding the evolution of migration strategies and the factors that shape individual decisions. Examining these dynamics across the full annual cycle is key to understanding how energy budgets influence survival and fitness.

Despite extensive migration ecology research, quantifying energy expenditure for small free-ranging migratory birds across their full annual cycles remains challenging. Their small body size limits the deployment of tracking devices that can simultaneously capture movement, behaviour, and environmental conditions over an entire year. Traditional methods (e.g., ringing or light-level geolocators) provide information on migration routes, stopover use, and wintering sites but cannot resolve fine-scale daily behavioural patterns and energy allocation across breeding, migration, and wintering periods.

Recent advances in multi-sensor data loggers now make it possible to integrate environmental (i.e., atmospheric pressure and ambient light) and biologging (i.e., movement and locomotion) data, enabling the reconstruction of hourly, daily, and annual energy budgets [11]. However, to date, few studies have attempted full annual reconstructions [12], leaving a major gap in our understanding of how energy expenditure and its allocation vary across the full annual cycle of migratory birds and among individuals exhibiting different migration strategies.

To quantify and compare energetic trade-offs among individual migratory birds, we developed an approach that integrates multi-sensor geolocator-derived activity, ambient light, and pressure data with environmental temperature records to reconstruct daily energy expenditure throughout the full annual cycle (see Fig 1). Energy use is partitioned into basal metabolism, locomotion, and thermoregulation, allowing detailed assessment of the costs associated with each migration strategy. Earlier studies have established fundamental methods for estimating field metabolic rates of birds during single annual phases [13] and across the annual cycle using time-activity budgets [14]. Building on these foundations, our framework extends this work by enabling reconstruction of complete annual energy budgets, capturing fine-scale variation in daily energy allocation during breeding, migration, and wintering periods. By incorporating historical climate data, it is also possible to evaluate how long-term warming trends have altered thermoregulatory demands at different site that birds visit and whether these changes have shifted the relative benefits of short-versus long-distance migration strategies. This combination of biologging, remote sensing, biophysical modelling, and climate analysis represents a novel framework for studying animal energetics of small migratory birds.

**Figure 1.**
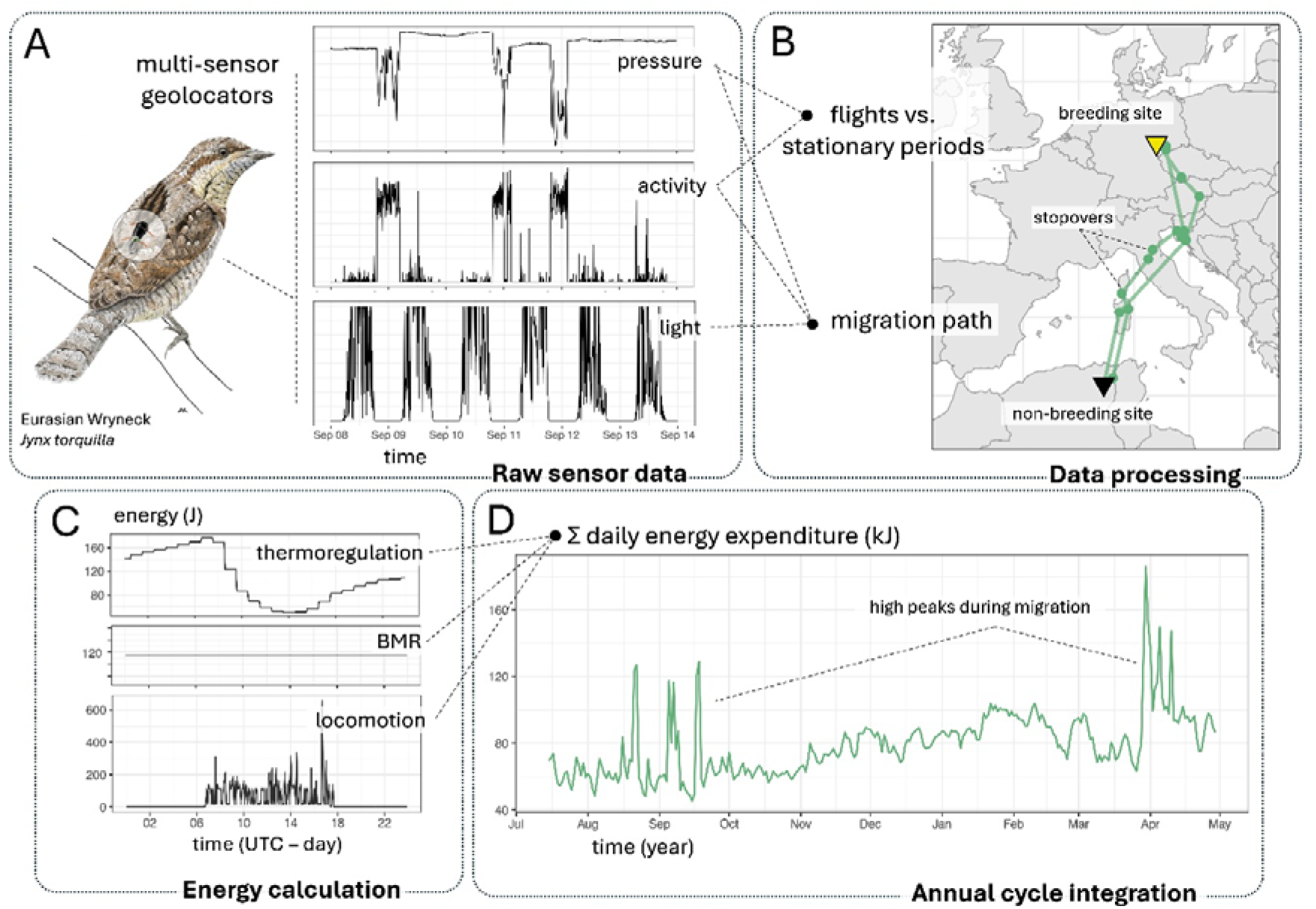
Reconstruction of annual energy budgets from multi-sensor logger data. (A) Eurasian Wrynecks were equipped with multi-sensor data loggers (geolocators) that harbour sensors for measuring atmospheric pressure, dynamic body acceleration (activity), and ambient light intensity at 5-min intervals year-round. (B) During the data processing, pressure and activity recordings are used to distinguish between migratory flights and stationary periods. Full migration tracks are reconstructed primarily based on time series of pressure recordings by matching them with global models of atmospheric pressure data and fine-tuning possible locations based on a global elevation model, light-level geolocation, and a movement model that implements the recorded flight durations with atmospheric conditions (wind) to obtain ground speed estimates during migratory flights (see methods for details). (C) Energy expenditure is estimated at 5-min resolution by combining three main sources of energy loss: (1) thermoregulation, based on remotely sensed temperature data along the migration track and species-specific thermoregulatory capacity, (2) basal metabolism (BMR), based on an allometric function developed for small-bodied birds, and (3) locomotion, based on accelerometer recordings and species-specific flight performance parameters. (D) Finally, annual energy budget is modelled as summed daily energy expenditure across the full annual cycle. Eurasian Wryneck (*Jynx torquilla*) illustration (c) Aline Knoblauch.

We applied this framework to data from multi-sensor geolocators deployed on Eurasian Wrynecks (*Jynx torquilla*), a small migratory landbird species, from a single Central European breeding population. The devices recorded atmospheric pressure, ambient light, and bird’s activity continuously [15] throughout the year, providing the detailed data necessary to estimate energy expenditure across breeding, migration, and wintering periods. Initial analysis of tracking data indicated substantial variation in migration distances among individual birds, with wintering locations ranging from southern Europe to northern and sub-Saharan Africa. This species model thus, provides a unique opportunity to examine how annual energy budgets vary among individuals that breed together but employ different migration strategies and overwinter in diverse climates.

Using this dataset, we addressed three main questions: (1) how does annual energy expenditure vary among birds originating from a single breeding population but employing different migration strategies; (2) which components of the energy budget (locomotion or thermoregulation) drive the variation; and (3) how has historical climate warming influenced thermoregulatory costs at wintering sites, potentially shifting the relative energetic advantage of alternative migration strategies. By answering these questions, we aim to illuminate the energetic trade-offs that shape migration decisions, demonstrate the utility of multi-sensor biologging for research of full annual cycle energetics, and provide a framework for predicting how migratory species may respond to the ongoing environmental changes.

## Results

### Migration strategy-specific energy budgets

We found an unexpected variation in migration distances and wintering destination for individual Wrynecks that breed near one another in Germany. Non-breeding sites spanned between Portugal (five tracks of four individuals, including one repeatedly tracked bird with IDs 28HV & 30BY in consecutive years), northern Africa (two individuals), and Sub-Saharan West Africa (two individuals; Fig 2a). Overall round-trip migration distances ranged between 4150 and 10650 km (mean ± SD: 6077 ± 2535 km). This large variation in migration distances also lead to variable length of autumn migration ranging between 13 to 91 days (35.3 ± 23.3 days). All birds resided at a single wintering site spending there between 142 and 206 days (186.1 ± 23.7 days). Spring migration was more concurrent lasting between 4 to 35 days (13.2 ± 9.5 days).

**Figure 2.**
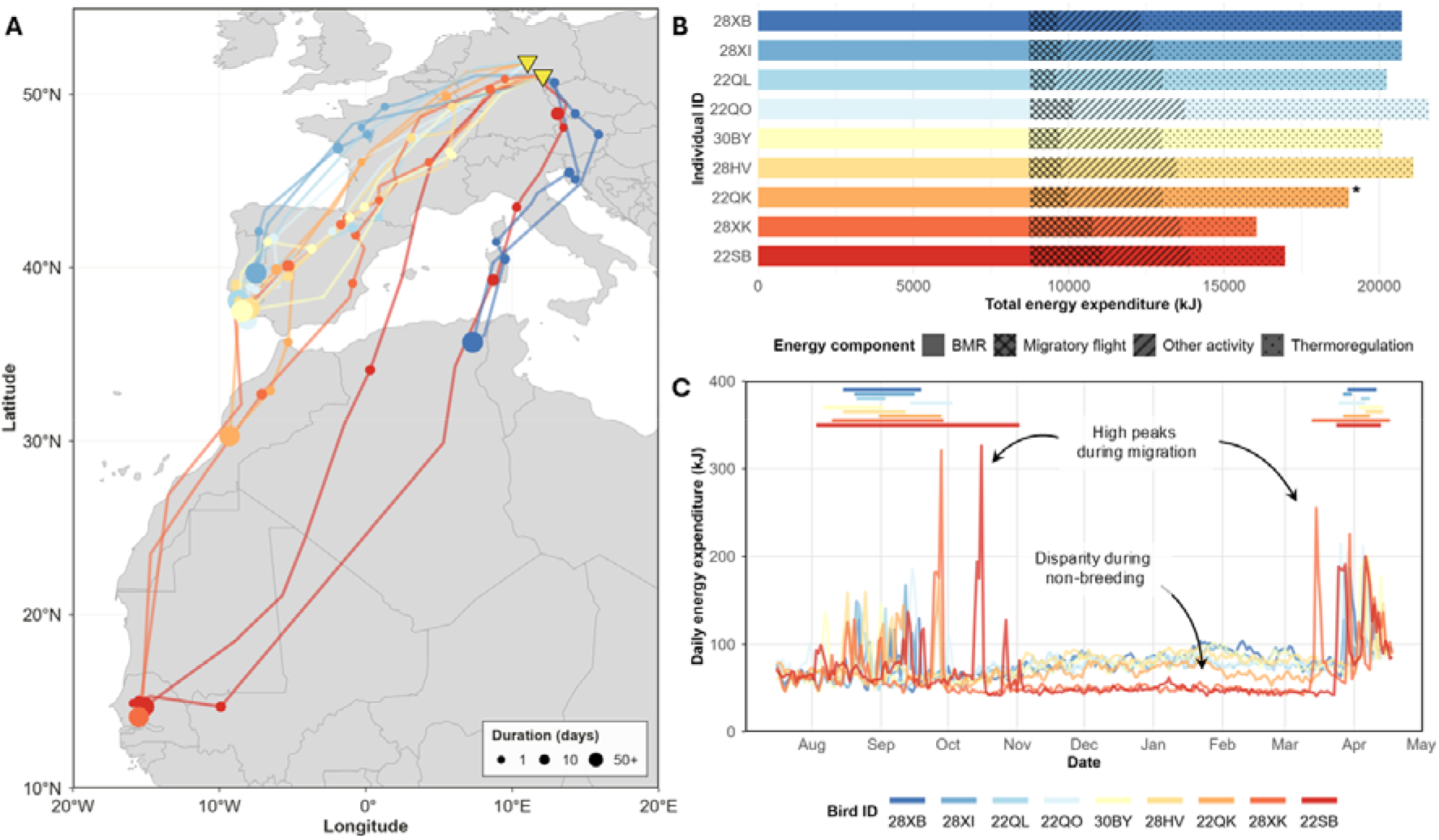
Annual migration patterns and energy expenditure of Wrynecks. (A) Wrynecks from the same breeding area in Germany (yellow triangles) show a large variation in migration strategies with non-breeding sites (large circles) spread from southwest Europe to northern and sub-Saharan Africa. Individual migration trajectories are shown in different colour arranged by migration distance from shortest (blue) to the longest (red). (B) Individuals employing different migration strategies differ in their estimated annual energy expenditure as well as the proportion of energy allocation to migratory flight, other locomotion costs, and thermoregulation, where longer migration distance is generally associated with lower annual energy expenditure. Bars represent summed energy budgets from 16-Jul to 8-Apr. Asterisk designates an individual whose accelerometer died before the start of spring migration and thus, migratory flight and locomotion costs for the missing 21 days were modelled based on autumn data (see methods). (C) Differences in annual energy budgets result from variation in daily energy allocation across the year, with contrasting migratory flight investment (peaks) and offset in thermoregulatory costs during the non-breeding period. Horizontal bars on top show autumn and spring migration periods from start to end.

Based on estimated locomotion, thermoregulation, and basal metabolism (BMR) costs, we found that outside the breeding period from mid-July until mid-April when birds are at different geographic locations, the average daily energy expenditure across all individuals was 73.1 ± 23.6 kJ day^−1^ (SD). This was portioned into 32.7 kJ (44.7%) for BMR, 24 ± 13.7 kJ (32.8%) for thermoregulation, and 16.5 ± 20 kJ (22.5%) for locomotion.

However, the energy allocation differed significantly based on migration distance: two long-distance migrants that crossed the Sahara invested 12.6–13.6% of their annual energy budgets for migratory flights but only 15.4–18% for thermoregulation, compared to 4–6.4% and 31.2–40.6% in other individuals, respectively. This led to over 25% variation in the total energy budgets among individuals outside the breeding season (Fig 2b) where longer migration distances were associated with lower annual energy expenditure (β = –0.715 ± 0.088 SE, t = –8.11, p < 0.001).

Daily energy expenditure showed clear peaks of high energy demand during migration (Fig 2c). Again, this was particularly pronounced for the two long-distance migrants for which the daily energy expenditure exceeding 9.5 times BMR on days when both individuals performed their 30+ hour long non-stop flights across the Sahara Desert, revealing extreme physiological stress that far surpasses demands of shorter distance strategies. This high energy demand, however, was offset by low thermoregulatory demands during the 5+ month long non-breeding residency period. A detailed comparison of annual energy expenditure at 5-min resolution of a long-distance migrant 28XK and a short-distance migrant 28XI Wrynecks is given in Figure 3.

**Figure 3.**
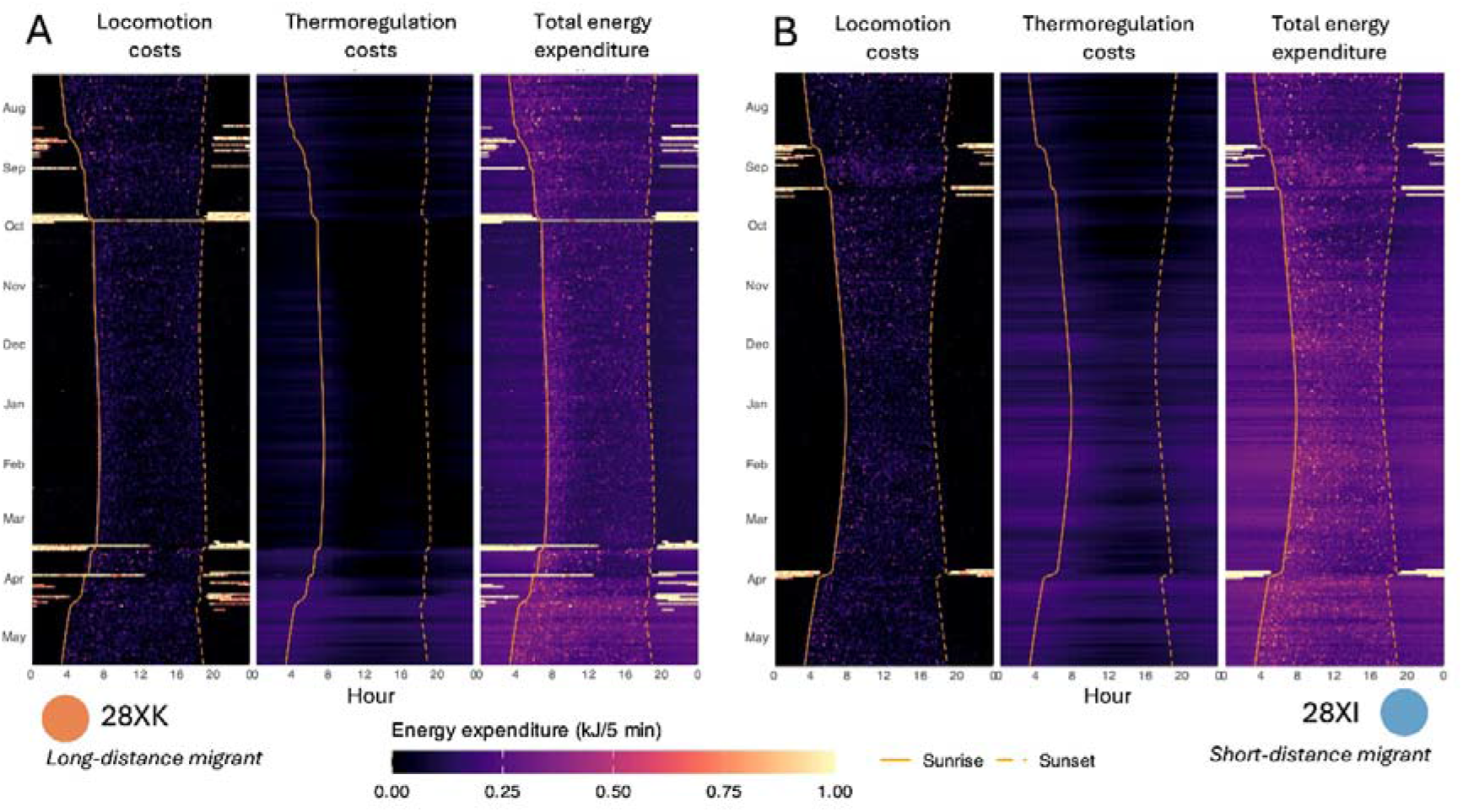
Annual energy expenditure of Wrynecks at 5-min resolution of (A) a long-distance migratory Wryneck (ID: 28XK) and (B) a short-distance migratory Wryneck (ID: 28XI). Energy expenditure is portioned into locomotion (left), thermoregulation (middle), and total energy expenditure including basal metabolism (right). Note the larger amount of high locomotory cost episodes corresponding to flight in the long-distance migrant while non-breeding thermoregulation costs from October to end of March are notably higher in the short-distance migrant. Total annual energy expenditure is markedly higher in the short-distance migrant amounting to a difference of 22.7% (19560 kj for LDM vs 24007 kJ for SDM).

### Winter thermoregulation under climate change

At all nine wintering sites identified though geolocator tracking, daily average temperature during the core non-breeding residency period between mid-October and mid-March has significantly increased since 1950 at an average rate of 0.018 ± 0.005°C^-year^ (Fig 4a). This has resulted in a significant decrease in daily over-winter thermoregulation costs at all sites at an average rate of 0.039 ± 0.015 kJ/day^-year^ (Fig 4b). Across the approx. 5-month long non-breeding period, this amounts to a substantial reduction in average thermoregulation costs by 450 ± 169 kJ over the 75-year period. However, this climate warming related thermoregulatory benefits are unevenly distributed depending on migration distance – winter period temperature increase has particularly benefited shorter-distance strategies where the decrease in winter thermoregulatory costs has been significantly faster (linear model with log-transformed distance; β_log(distance)_ = 0.032 ± 0.008, t = 4.13, p = 0.004).

**Figure 4.**
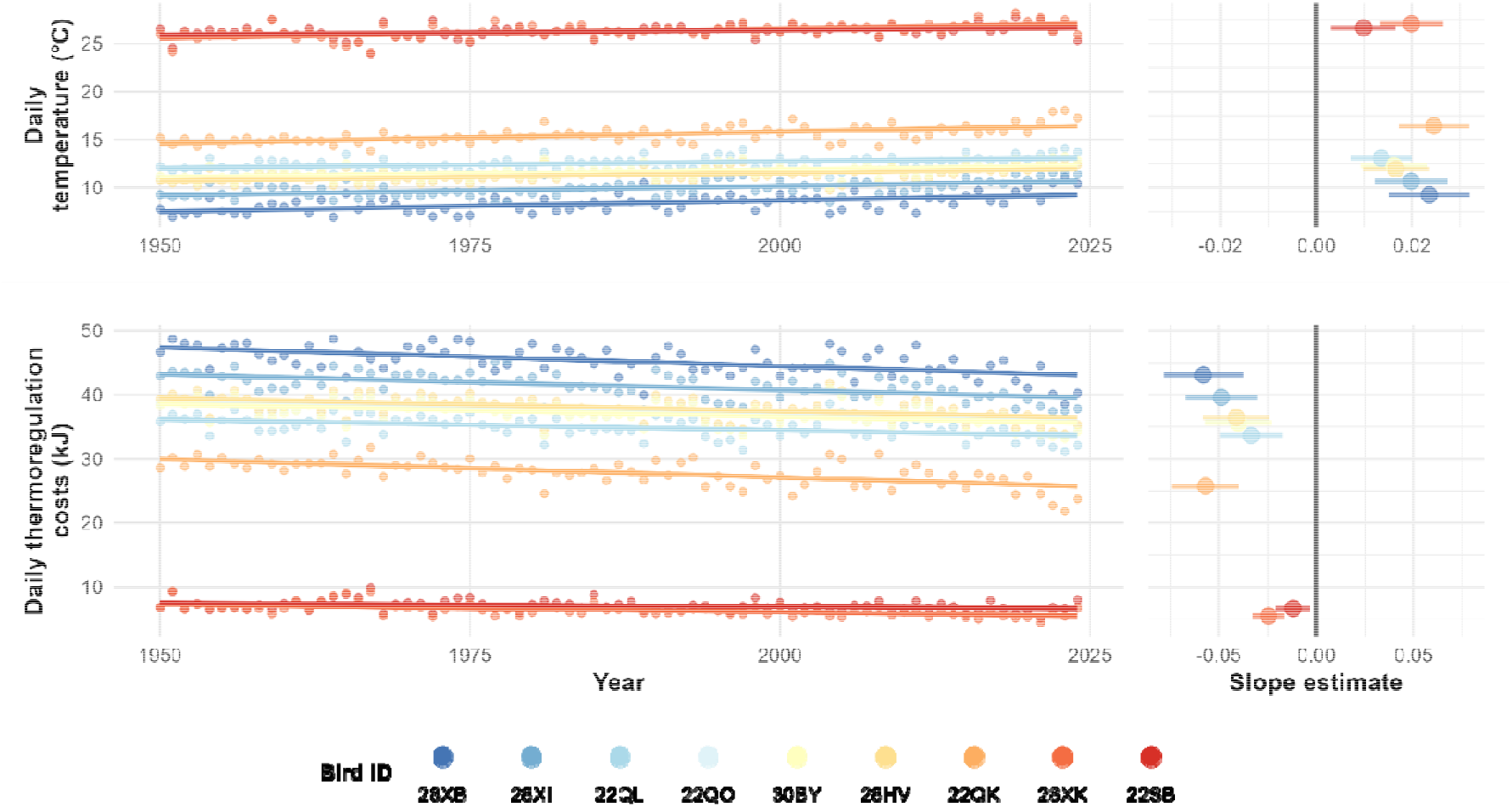
Changes in average daily (A) temperature and (B) estimated thermoregulation costs from 1950 to 2025 of nine geolocator-tracks of Eurasian Wrynecks at their respective wintering sites between 15-Oct and 15-Mar. Left panels show annual values (points) with fitted least-squares regression lines. Right panels display slope estimates and their 95% CIs from the regression analyses.

## Discussion

Migratory birds need to balance energetic trade-offs across the annual cycle where a costly investment in long-distance migratory movements may pay off through lowered thermoregulation costs. Yet, tracking and quantifying year-round energy expenditure of small-bodied migratory birds in the wild has been challenging as they often move thousands of kilometres between breeding, stopover, and wintering locations [12]. Because of this, empirical estimates of full annual energy expenditure have been largely restricted to large species for which relatively heavy multi-sensor transmitters are suitable [5,16], leaving the vast majority of smaller migrants effectively understudied in this respect. Here, we present a framework that enables achieving this feat at a high spatiotemporal resolution through integration of multi-sensor biologging and biophysical models of thermoregulation and locomotion costs.

The acquired average energy expenditure estimate of 73.1 kJ day^−1^ for Wrynecks, fall within the expected range based on doubly labelled water method-derived allometric relationship, according to which, an approx. 36 g sized bird should have a daily energy expenditure of 70–130 kJ, depending on season and activity level [17]. The peak daily energy expenditure of the tracked long-distance migrant Wrynecks exceeded 9.5 times BMR or >300 kJ day^−1^, which is at the upper extreme of the expected daily energy expenditure capacity of birds [18]. In our dataset, these extreme values were recorded only for long-distance migrants that both performed more than 30-hour long non-stop flights to cross the Sahara Desert, thus, spending 100% of the day in flapping flight.

Some sources of uncertainty remain in the presented approach for the estimation of energy budgets limiting the precision of absolute estimates. For example, microclimates that birds inhabit can affect thermoregulation costs and the used biophysical models does not account for costly tissue syntheses like feather moult. As multi-sensor biologging is becoming a widely used tool in bird migration research over the last decade [19], this opens new opportunities to reanalyse the existing data or be applied to entirely new dataset with wide-ranging ecological question. Moreover, because our framework does not rely on long-distance movements per se, but on linking behaviour-specific energy costs, it can be readily applied to resident species as well as to shorter-term studies targeting specific ecological questions (e.g., reproductive investment and success, carry-over effects, season-, habitat- or weather-dependent variation in energy expenditure, etc.).

### Intraspecific variation in migration strategies and energy budgets

Our tracking data revealed striking intraspecific variation in migratory behaviour among Eurasian Wrynecks from the same Central European breeding population. Individuals exhibited large differences in both migratory routes and wintering destinations: some travelled via the Apennine Peninsula, others via the Iberian Peninsula, with wintering areas ranging from southern Portugal to sub-Saharan West Africa. To our knowledge, this represents one of the most pronounced cases of migratory strategy polymorphism documented in a small landbird (see also [20]).

In many songbirds, migratory phenotypes tend to follow relatively clear geographic structuring among populations, particularly in terms of migration distance and orientation [21]. Crossbreeding experiments in songbirds have demonstrated that migratory orientation and distance are heritable quantitative traits [22,23]. The persistence of short-and long-distance migrants within a single local breeding population therefore raises interesting questions about how migratory programs are encoded in this species and about selective pressures and mechanisms that maintain this diversity. Because the Eurasian Wryneck (Piciformes) shares its most recent common ancestor with Passeriformes dating back around 80 million years [24,25], it is plausible that the mechanisms underlying migratory program encoding in this species differ from those described in passerines [26,27].

Regardless of the underlying encoding mechanisms, the co-existence of multiple migratory strategies may reflect a stable polymorphism maintained by frequency-dependent or condition-dependent selection. Under such scenarios, the relative fitness payoffs of different strategies may fluctuate across years depending on environmental conditions experienced across the annual cycle. Yet, our energetic analysis adds an important dimension to this picture. The fact that shorter migrations incur substantially higher total non-breeding energy expenditure, yet they persist (and dominate) within the population, suggests that energetic costs alone do not determine the adaptive value of a given strategy.

A recent tracking study from Switzerland identified short-distance migration as the dominant strategy of Wrynecks from this part of the breeding range [28] which came as a surprise since all northern and central European populations were thought to be long-distance migrants wintering in Sahelian Africa [29,30]. This, in combination with indications of increasing wintering population on the Iberian Peninsula [31], may suggest rapid evolutionary changes in Wryneck’s migratory behaviour over the last several decades. In this light, the co-existence of multiple migratory strategies within our population may represent a transitional stage on the evolutionary timescale, as migration distances shrink and a shift from long- to short-distance migration strategies unfolds.

While from a purely energetic perspective, shorter distance strategies may not be favoured, shorter migrations may offer multiple additional benefits that could outweigh the higher annual thermoregulation needs. First, birds wintering closer to their breeding grounds tend to return earlier in the spring [32], a pattern commonly associated with higher reproductive success [33]. This timing advantage can be critical for coping with climate change induced phenological shifts in the lower trophic levels [34,35]. Second, long-distance migration and especially crossing of ecological barriers (e.g., the Sahara Desert) is associated with higher mortality rates [4]. This can result from adverse weather and harsh environmental conditions encountered *en route*, or extreme physiological and energetic demands during long-distance endurance flights that we observed in tracked Wrynecks. Disentangling the relative contributions of these factors to lifetime fitness will require multi-year tracking combined with migration strategy-specific data on reproductive success and mortality.

### Shifting energy budgets under climate change

Thermoregulation emerged as the dominant driver for the variation of non-breeding energy expenditure across individuals, accounting for approx. 1/3 of the total non-breeding energy budget in short-distance migrants, compared to just 16.7% in long-distance migrants. This finding has important implications when viewed in the context of the ongoing climate change, as rising winter temperatures at higher-latitude wintering sites has already and will continue to disproportionately reduce the thermoregulatory burden faced by short-distance migrants.

Shortening of migration distances is generally expected with rising temperatures and has been documented in several species [36,37]. However, this phenomenon is mostly observed for short-distance and partially migratory species where gradual shortening of migration distances or increased propensity for residency is observed. For traditional long-distance migrants, a gradual shortening of migration distance is more difficult to achieve because the Sahara Desert represents an inhospitable ecological barrier along the way and thus, it requires a big jump[Z]like reduction in migration distance [38]. Since migration distance is at least partially genetically encoded [39], environmental conditions need to reach a critical turning point that enables and promotes such jump[Z]like shift. A significant reduction in thermoregulatory needs found in our study, represents one such aspect, which alongside ca. 50% lower migratory flight costs and potential reproductive/survival implications mentioned above, is likely contributing towards an increased preference for shorter migration distances and, hence, a shift from long- to short-distance migration in Central European Wrynecks.

Despite the continuous reduction of thermoregulatory needs over the last 75 years, shorter migrations are still associated with 20-25% higher annual energy demands compared to trans-Saharan migration. Conversely, Linek et al. [8] found similar disparity in winter thermoregulation costs as in our study but an overall annual energetic equivalence for short-distance migratory and resident Common Blackbirds *Turdus merula*. These two examples provide contrasting results for how annual energy budgets differs among individuals from the same breeding population but contrasting migration strategies and wintering areas. Alternative migration strategies undoubtedly entail distinct trade-offs between energetic costs, mortality risks, and fitness which will continue to transform and adapt as environmental conditions continue to change. Mechanistic migration models that couple physiology, energetics, demography, and changing environmental conditions highlight that resilience to global and climate change is strongly strategy-dependent, but their predictive power hinges on robust empirical characterization of how individuals and populations use space, time, and energy throughout the annual cycle [40]. Hence, identifying population-specific migration strategies and understanding the associated energetic, mortality, and fitness consequences are essential for predicting how migratory species will respond to the ongoing global changes.

## Materials and methods

### Tracking

We conducted a tracking study on Eurasian Wrynecks at two nearby breeding sites in northeast Germany (approximately 51.82°N, 11.05°E and 51.1°N, 12.1°E). At both locations, wrynecks breed in provided nest boxes, and we captured adult birds using mist nets during chick feeding. Captured wrynecks were fitted with GDL-PAM3.0 multi-sensor geolocators (manufacturer: Swiss Ornithological Institute) which measured ambient light, atmospheric pressure, and dynamic body acceleration every 5 minutes. Acceleration measurements were taken along the z-axis for 3.2 seconds at 10 Hz frequency, with the geolocators pre-processing this data on-board and storing the sum of absolute differences between consecutive measurements (3.2 sec at 10 Hz = 32 consecutive measurements).

In total, we tagged 40 individuals in 2019, 45 in 2021, and 35 in 2022. The average mass of a geolocator was 1.4 ± 0.05 g (SD) while the average body mass of the tagged birds was 36.0 ± 1.9 g (n = 35). Since geolocators do not transmit data, they need to be recovered for data download. In subsequent years, we recovered 9 out of the 120 devices deployed, a recovery rate of 7.5%. The low overall recovery rate is likely explained by the Eight of the nine recovered geolocators contained full annual cycle data. Due to memory shortage, one device (id: 22QK) stopped recording acceleration on 18 March shortly before the start of spring migration, resulting in incomplete activity recordings (pressure and light recordings lasted until 10 April and covered full spring migration).

Tracking data was analysed using R-package *GeoPressureR* (v.3.2.0) following standard procedures outlined in Nussbaumer et al. [41,42]. In short, *GeoPressureR* matches pressure data recorded by the loggers to global weather reference data (ERA5 hourly surface level pressure dataset) to construct a likelihood map of positions for each stationary period where there is a match in the ground level elevation, considering the temporal variation and likelihood of the mismatch. Migratory flights were manually distinguished from stationary periods based on simultaneous drops in pressure and sustained high dynamic body acceleration using the TRAINSET tool (https://trainset.raphaelnussbaumer.com/). Light data was used to create likelihood maps for each twilight using the threshold method [43], further refining the position estimates from pressure. Finally, we computed transition probabilities between stationary positions by applying a movement model that integrates birds’ airspeed and wind data (ERA5 hourly data on pressure levels) at the time and respective altitude of each flight. Thus, the resulting modelled path from *GeoPressureR* is the path that maximizes the overall probability between stationary sites from the pressure and light maps, accounting also for the ground speed distribution of the bird.

### Estimating energy expenditure

We calculated daily energy expenditure as a sum of three main sources of energy loss: (1) basal/resting metabolism (BMR), (2) thermoregulation, and (3) activity, excluding energy loss associated with digestion, energy retention (storage), and tissue synthesis.

#### Basal metabolism

BMR in birds has been shown to scale with body mass [44] and thus, can be estimated applying an allometric function. We calculated BMR (mlO_2_/h) of wrynecks using the allometric relationship described across 211 avian species [45] as BMR = 6.7141M^0.6452^, where M corresponds to wryneck body mass in grams = 36.0 g (average body mass of individuals in our study). We then converted the BMR measurement into kJ per day using the conversion factor of 20.1 kJ/l O_2_ [46].

#### Thermoregulation costs

Earlier studies have indicated that endotherms have a thermoneutral zone wherein thermoregulation costs are 0, but outside this zone, costs increase linearly [47]. Similar to BMR, thermoregulation costs in birds outside of the thermoneutral zone have been shown to scale with body mass [45].

The boundaries of the thermoneutral zone are delineated by the lower critical temperature (T_LC_) and the upper critical temperature (T_UC_). Following the Scholander–Irving model of heat transfer [48] the lower critical temperature for resting endotherms is described as T_LC_ = T_B_ − BMR/C, where T_B_ represents body temperature (40°C for avian species), BMR denotes basal metabolic rate in mlO_2_/h, and C signifies the rate of heat loss or thermal conductance. Analogous to BMR, thermal conductance (C) in avian species has been demonstrated to scale with body mass and based on empirical measurements across 211 avian species, this relationship can be described using the following allometric equation: C = 0.8248M^0.5088^, where M corresponds to body mass in grams [45]. Following this, thermoregulation costs (TC) at ambient temperatures below T_LC_ can be calculated as TC = (T_LC_ - T_A_)/C, where T_A_ corresponds to ambient temperature.

Thermoregulation costs of birds at ambient temperatures above T_UC_ (i.e., costs of cooling) have been poorly quantified previously but follow a similar linear increase as costs of heating at ambient temperatures below T_LC_ [49]. Thus, we assumed the same linear increase in thermoregulation costs at ambient temperatures above T_UC_: TC = (T_A_ – T_UC_)/C, where T_UC_ = 34.3°C corresponding to the mean T_UC_ as quantified across 167 avian species [50].

We obtained hourly ambient temperature data experience by individual wrynecks across the full annual cycle along their migration tracks (as estimated using *GeoPressureR*, see above) from ERA5-Land hourly data dataset (or ERA5 hourly data on single levels dataset when the birds were in flight over sea/ocean) [51]. To this end we used air temperature at 2m above the surface at hourly resolution and converted the thermoregulation costs into kJ per hour using the conversion factor 1 J/s = 172 mlO_2_/h. Likewise, we obtained annual historic temperature records dating back to 1950 along the migration routes of geolocator-tracked Wrynecks from the ERA5 dataset.

#### Activity costs

Flight costs in birds can be estimated using the aerodynamic power model [52]. We calculated the metabolic power of an average Wryneck using the animal flight performance tool from the R-package *afpt* [53]. Apart from the body mass (we used the average of our study birds = 36.0 g), other morphological parameters (wingspan and wing area) for the model were obtained from [54]. We assumed that during migration birds followed a time minimization strategy [3], and consequently we used *afpt* function *findMinimumTimeSpeed* to estimate flight costs of wrynecks. Adopting this, energy expenditure of an average flight was estimated at 196.2 J/min.

We then calculated the average activity as recorded by the geolocator’s accelerometer during migratory flights across all tracked Wrynecks (mean = 35 ± 8 SD, n = 11724 recordings). The activity measurement of 35 was then assumed to be flight under normal conditions and thus correspond to energy expenditure of 196.2 J/min as estimated with the animal flight performance tool (see above).

Overall, the measured activity levels ranged between 0–75 across all individuals and all recorded behaviours (including non-migratory flight), where 0 corresponds to no movement and thus, no energetic costs of locomotion. Previous studies have shown that energetic costs of basic behaviours with minimal body movement, like alert perching, roughly correspond to energetic costs of BMR [49]. Thus, we assigned an activity measurement of 1 to correspond to the energy expenditure equal to BMR. Further, we used these two known measurements (activity level 1 = BMR = 22.71 J/min and activity level 35 = flight = 196.2 J/min) as anchor points for linear extrapolation of energetic costs of all other activity level measurements.

For one track (ID: 22QK), acceleration measurements were unavailable during the final 21 days of spring migration because the accelerometer stopped recording on 18 March. Pressure and light data remained available and were used to identify stationary periods and migratory flights. Locomotion costs during the missing period were reconstructed using locomotion costs derived from the same individual’s autumn migration period that was randomly sampled each 5-min for either migratory flight costs or other locomotion costs, i.e., at stopover sites. The latter was done separately for day and night since birds are mostly inactive during nighttime.

### Statistical analyses

To test whether annual energy expenditure varied with migration distance, we fitted a linear model with total energy expenditure (kJ), summed over the period from 16 July to 8 April, as the response variable and total migration distance (km) as the predictor. This analysis comprised nine annual tracks from eight individuals because one individual was tracked in two consecutive years under different logger IDs. These two tracks were retained as separate observations because they represented energy expenditure under different annual environmental conditions and individual behaviours.

To quantify historical changes in winter conditions, we calculated mean daily ambient temperature and mean daily thermoregulation costs during the core wintering period from 15 October to 15 March for every year between 1950 and 2025. Calculations were performed separately for each of the nine tracking-derived wintering locations. For each wintering location, we fitted separate linear regressions of mean daily temperature and mean daily thermoregulation cost against year. We extracted the temporal slope, its standard error, test statistic, p-value, and 95% confidence interval from each model.

To test whether the climate-associated changes in winter thermoregulation costs varied with migration distance, we used the track-specific temporal slopes from the thermoregulation models as the response variable. We fitted an inverse-variance weighted linear regression in which the natural logarithm of total migration distance was the predictor and each slope estimate was weighted by the inverse of its sampling variance, 1/SE^2. This gave greater influence on temporal slopes estimated with higher precision. Migration distance was log-transformed to represent proportional differences in distance and to reduce the influence of the two long-distance migration tracks

All data analyses were performed in R 4.5.0. Descriptive values are reported as means ± SD, whereas model coefficients are reported as estimates ± SE. All statistical tests were two-sided, and effects were considered statistically significant at p<0.05. Model-derived uncertainty is presented using 95% confidence intervals.

## Acknowledgements

We thank Bill Buttemer and Felix Liechti for thermoregulation and flight cost discussions.

